# Inactivation of Microorganisms in the Complex Regions of Transvaginal Ultrasound Probes By a UVC-LED Light Based Disinfection System

**DOI:** 10.64898/2026.03.23.713795

**Authors:** Muhammad Yasir, Mark D.P. Willcox

## Abstract

Endocavity ultrasound transducers, particularly transvaginal ultrasound (TVUS) probes, contain intricate structures such as notches, grooves, lens surfaces, and handle edges that are highly susceptible to microbial contamination yet difficult to disinfect using conventional high-level disinfection (HLD) methods. This study evaluated the efficacy of a novel ultraviolet-C light-emitting diode (UV-C LED) HLD system in eliminating microbial contamination from these complex probe surfaces. Two TVUS probes were sampled from predefined high-risk regions before and after disinfection following clinical use. Probe A was sampled at the top and bottom notches and both sides of the handle, while Probe B was assessed at the lens, edges, and bent groove regions. Microbial contamination was quantified using swab sampling, culture on agar plates, and identification via MALDI-TOF. Environmental sampling of examination and disinfection rooms was also performed. To assess this system robustness, probe sites were repeatedly inoculated with *Bacillus subtilis* spores and evaluated following UV-C treatment. Before UV-C treatment, contamination rates ranged from 25% to 57% across sampled regions, with microbial loads reaching up to 3.9 log₁₀ CFU. Identified organisms included *Staphylococcus epidermidis*, *Pseudomonas koreensis*, *Bacillus cereus*, and *Propionibacterium* spp. Probe sheaths were also predominantly contaminated with *Staphylococcus epidermidis.*, with counts reaching up to 4.3 log₁₀ CFU, Environmental sampling revealed diverse microbiota, with higher contamination levels in examination rooms compared to disinfection areas. Following 90 seconds of UV-C exposure, no microbial growth was detected on any sampled site, indicating 100% decontamination. Additionally, UV-C treatment achieved >6.7 log₁₀ reduction of *B. subtilis* spores across all tested regions. These findings demonstrate that UV-C LED technology provides rapid, effective, and consistent high-level disinfection of complex TVUS probe surfaces, supporting its potential as a rapid and reliable disinfection modality in clinical setting.

## Introduction

Ultrasound transducers (also known as probes) are used for a variety of diagnostic and procedural applications, including transvaginal (TV), transrectal (TR), transoesophageal echocardiography (TOE), and imaging of open wounds. During these procedures, the transducer may come into contact with mucous membranes of the vagina, rectum, oropharynx and oesophagus, as well as non-intact skin such as ulcers or open wounds. Importantly, when they are used in sterile body areas, such as the intravascular space, there might be high chances of risk of infection. Certain structural features of ultrasound probes, including notches, grooves, seams, and textured surfaces, may increase the risk of contamination. These areas can retain biological material or microorganisms, if not properly disinfected.

Inadequate reprocessing of transvaginal ultrasound (TVS) probes can lead to cross-contamination and nosocomial infections, posing a serious risk to patients’ health, in particular to pregnant patients. (1, 2). Commonly, the vaginal microbiota such as *Gardnerella vaginalis, Ureaplasma urealyticum, Candida albicans, Prevotella* spp., and *Lactobacillus* spp have been found associated with probes infection (3, 4). Pathogens not usually associated with the normal microbiota of the healthy vagina including human papillomavirus (5), Epstein–Barr virus 9 (6), human immuno deficiency virus (HIV), cytomegalovirus, Gram-negative bacteria such as *Escherichia coli*, *Pseudomonas aeruginosa* and Klebsiella sp, Gram-positive bacteria such as methicillin-resistant *S.aureus*, *Clostridium difficile,* vancomycin-resistant Enterococcus spp, *Neisseria gonorrhoea, Treponema palladium*, and *Mycobacterium avium* can also contaminate the probes and may cause disease (7).

Probe contamination may also occur due to perforation, tearing, or leakage of the probe cover during use, allowing body fluids or microorganisms to contact the probe surface. This can account for up to 9.0% of infections (8–10). Reported leakage rates range from 0.4% to 13% for condoms and 0% to 5% for commercial covers (11). Additionally, there is an estimated 9% risk of condom perforation in patients undergoing transrectal biopsy under ultrasound guidance (10). Similarly, perforation rates of up to 81% have been reported for commercial probe covers, resulting in contamination of transducers with blood and other body fluids (8, 10).

Most ultrasound transducers cannot undergo established sterilisation processes; therefore, it is imperative that correct high level disinfection (HLD) is undertaken. In a previous study (6), UVC based Lumicare HLD disinfectant system disinfected transvaginal ultrasound probes (TVUS) within 90 seconds and effectively eliminated microorganisms associated with them. However, to reinforce its effectiveness, the current study focussed on the most challenging probe areas (notches, seams, grooves, and textured surfaces), commonly referred to as potential cold spots of the probes (cold spots being defined as areas where light, including ultraviolet light may not easily reach). Therefore, the current study assessed the effectiveness of a UVC system in reducing microbial load in the complex areas of two different types of ultrasound probes.

## Methods

### Ex vivo testing

This study was approved by the Human Research Ethics Committee of the University of New South Wales (HREC number: HC220136). The ultrasound probes were collected after use in an in vitro fertilization clinic (Connect IVF, Sydney, Australia) from February 26 to April 02, 2025. These had been sheathed with a non-sterile latex free probe cover (Sonologic) prior to use. After use, the sheath was removed, and sterile cotton swabs (Bacto Laboratories, Mount Pritchard, NSW, Australia) were passed twice over a 2-5 cm^2^ area from the probes including from the top notched, bottom notched, left side of the handle, right side of the handle of one probe, grooves (bent areas) and lense areas of other probe (randomly selected) to examine the numbers and types of microbes present on the endoscopes prior to disinfection (Figure 1).

**Figure 1:**
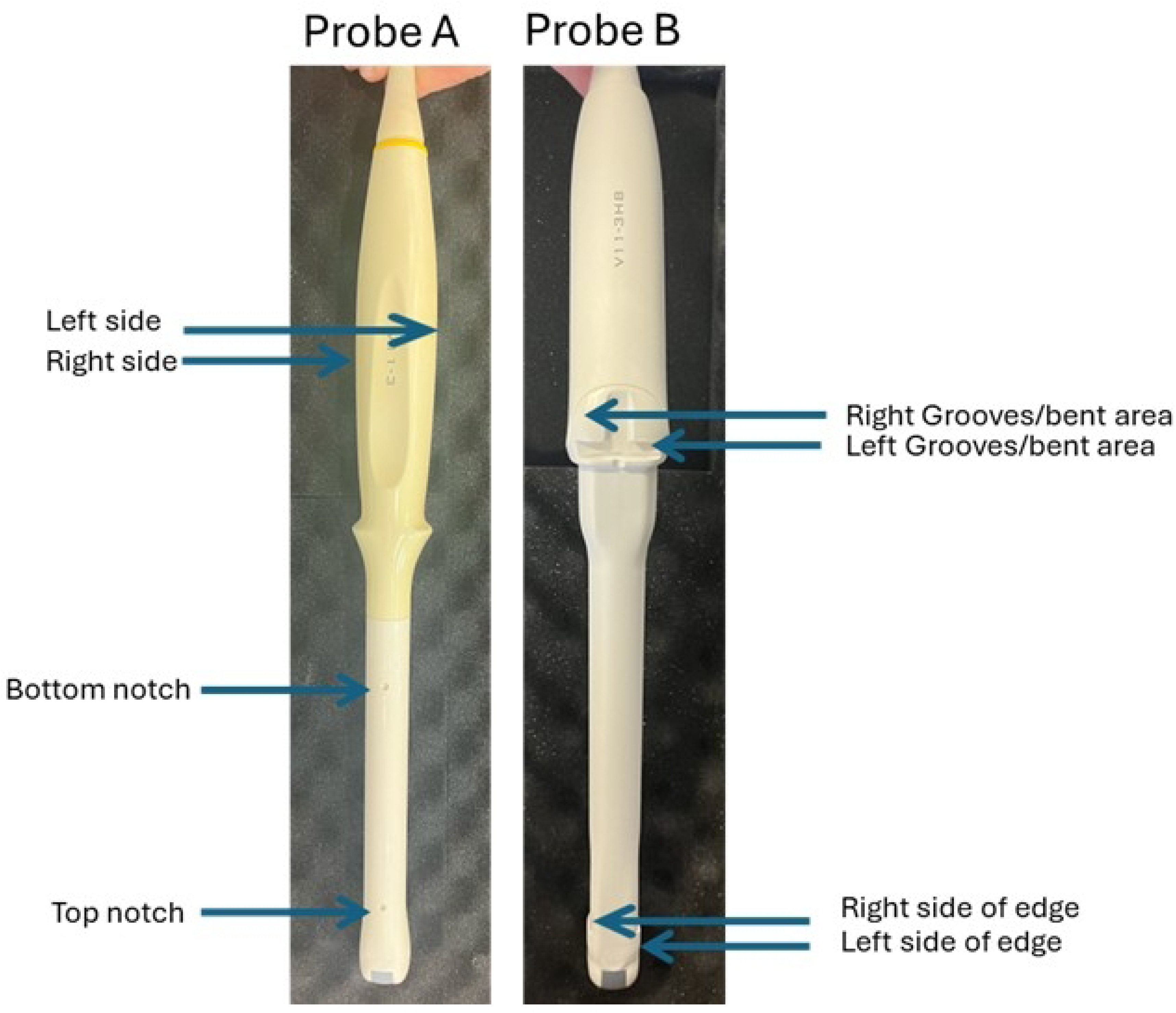
Ultrasound probes used in the current study. The blue arrow pointed the areas from where samples were swabbed before and after UVC treatment for isolation of microorganisms.

The ultrasound probes were then loaded into the Lumicare ONE High Level Disinfection UVC system (Lumicare 606A743U7Y, Sydney, Australia) and the system run for a standard time of 90 seconds. After disinfection, the ultrasound probe was removed, and the opposite side of each probes’ sites was swabbed as described above. The swabs were then put into AMIES transport media (Bacto Laboratories, Australia) and transported to the microbiology laboratory of the School of Optometry and Vision Science University of New South Wales (UNSW) in cold chain and processed within 4 hours of collection.

To assess whether the sheaths (protective covering of ultrasound probes) had any bacterial contaminations, five sheaths were randomly selected and analysed for the presence of contaminants. The sheaths were swabbed from the internal surfaces over a 5-10 cm^2^ area which had directly contacted the probes.

All the swabs were processed under aseptic conditions. Microorganisms were recovered in Phosphate Buffer Saline (PBS; NaCl 8 g/L, KCl 0.2 g/L, Na_2_HPO_4_ 1.4 g/L, KH_2_PO_4_ 0.24 g/L; pH 7.4) by vortexing the swabs in vials in the presence of glass beads (0.5 mm diameter, Sigma Aldrich St Louis, MO, USA) for 3-5 minutes. 200 μL of the resulting slurry was spotted onto chocolate blood agar (CBA; Beckton Dekickinson, Maryland, USA) and incubated in three different atmospheric conditions (aerobic, microaerophilic and anaerobic) for 24-72 hours at 37°C. 200 μL was also plated onto Sabouraud’s agar (SDA; Beckton Dekickinson, Maryland, USA) and incubated for 5-7 days at 25°C to grow any fungi present on the endoscopes. Following incubation, numbers of similar colonies of each bacterium and fungus were counted and purified. Pure cultures of microbes were stored in 25% glycerol at −80 °C.

#### Environmental sampling

To assess whether there were any microorganisms (bacteria, fungi, yeasts) present in the environment during the process of sampling and disinfection, environmental monitoring was assessed using agar plates. One plate of each CBA and SDA with opened lids were placed in the clinical and disinfectant rooms for the entire duration of sample collection and disinfected in the Lumicare device.

#### Microbial identification

Microbes were recultured on CBA and incubated overnight. A loopful of growth was suspended in 300 μL of HPLC-grade water, mixed, and treated with 900 μL absolute ethanol. After centrifugation (13,000 rpm, 2 min) and drying, pellets were dissolved in 25 μL of 70% formic acid and 25 μL acetonitrile, then centrifuged again. One μL of the supernatant was spotted on a MALDI plate, overlaid with HCCA matrix, air-dried, and analysed by MALDI-TOF for microbial identification (12).

### *In vitro* testing Sterilising the probes

To confirm that the ultrasound probes used in this study were sterile at the start and after every disinfection cycle, they were cleaned with 4% bleach for 4–5 minutes before each *Bacillus* spore inoculation and then dried for 20–30 minutes. After decontamination with bleach, the probes were rinsed with sterile Milli-Q water to remove any residual bleach and prevent interference in subsequent use. Thereafter, each probe was swabbed and the swab suspended in PBS within a sterile test tube and mixed by vortex for 3-5 minutes at ambient temperature (21 °C). The suspension was further diluted in the same diluent. Aliquots of 1 mL were plated onto Tryptic soya agar plates (TSA; Oxoid, Basingstoke, UK) and incubated for 24 h at 37 °C. After incubation, the agar plates were examined for bacterial growth.

Two ultrasound probes (Mindray, China; Figure 1) with notches at the top and middle, one bent at the centre were disinfected with 4% ethanol and dried before use.

#### Spore production

The spores of *Bacillus subtilis* ATCC 19659 were produced following a published method (6). Then spores in suspension (1×10^9^ spores/mL; 20 μL) were dried for 30–45 min on probe notches, bends, and transducer lenses. Probes were then either left at room temperature (21°C) or exposed to UVC for 90 sec using the Lumicare ONE HLD system (S.N: 6LMEC6YEUA, 20–25°C). Following treatment, spores were recovered by swabbing the inoculated areas. Swabs were vortexed in PBS containing glass beads for 3–5 minutes, followed by serial dilution and plating onto tryptic soy agar (TSA). Plates were incubated at 37 °C for 24 h. Similarly, spores were recovered from control areas of the same surface that did not receive UVC treatment. Uninoculated areas were processed using the same procedure to confirm sterility. No neutralising agent was required, as UVC exposure does not result in residual antimicrobial activity.

### Statistical Analysis

The numbers and type of microbes isolated from the transvaginal ultrasound probes after use and then after a disinfection cycle in the Lumicare ONE High Level Disinfection device was compared using nonparametric (Mann Whitney test). Recovery of *Bacillus* spores before and with UVC treatment from different sites of probes was compared using t test with Graph Pad Prism 8 version 8.0.2. A P value less than 0.05 indicated statistical significance.

## Results

### Ex vivo testing

Areas of probes A and B were heavily contaminated with microbes before UVC treatment. A diverse range of bacterial species was detected on each probe. For probe A, these included Staphylococcus, Pseudomonas, Propionibacterium, Bacillus, Legionella, and Tsukamurella spp. Nearly all sites of the probe were contaminated with multiple bacterial species (Table 1).

**Table 1.** Numbers of identified bacteria on different sites of probes A and B before treatment with UVC.

| Patient ID | Top Notch untreated<br>(Log<sub>10</sub> CFU) | Bottom Notch<br>untreated<br>(Log<sub>10</sub> CFU) | Right side of<br>handle<br>untreated<br>(Log<sub>10</sub> CFU) | Left side of handle<br>untreated<br>(Log<sub>10</sub> CFU) |
| --- | --- | --- | --- | --- |
| **Probe A** |  |  |  |  |
| **Sample 1** | • *S. epidermidis* (1.6)<br>• *Pseudomonas fulva* (1.6) | Nil | Nil | Nil |
| **Sample 2** | • *Propionibacterium sp* (3.5) | Nil | Nil | Nil |
| **Sample 3** | • *Pseudomonas fulva* (3.5) | Nil | Nil | Nil |
| **Sample 5** | Nil | Nil | *S.warneri* (2.6)<br>*S.lugdunensis* (2.3) | Nil |
| **Sample 6** | Nil | Nil | *S.warneri* (2.9)<br>*Legionella pneumophila* (1.8) | Nil |
| **Sample 9** | *S. epidermidis* (2.2) | *S. epidermidis* (2.7) | Nil | *Pseudomonas balearica* (3.1)<br>*Bacillus amyloliquefaciens* (3.7) |
| **Sample 10** | Nil | *S. epidermidis* (2.7) | Nil |  |
| **Sample 15** | Nil | Nil | *Tsukamurella hongkongensis* (3.1) | Nil |
| **Probe B** |  |  |  |  |
| **Sample 17** | *Legionella pneumophila* (2.8) | Nil | Nil | Nil |
| **Sample 18** | *B. cereus* (3.6) | Nil | *Pseudomonas koreensis* (3.8) | Nil |
| **Sample 19** | Nil | Nil | *S. epidermidis*(1.8) | Nil |
| **Sample 20** | *B. cereus* (2.1) | Nil | *S. epidermidis*(3.3) | Nil |
| **Sample 22** | Nil | Nil | *Bacillus infantis* (3.5) | Nil |
| **Sample 23** | Nil | Nil | Nil | *S. capitis* (1.9) |
| **Sample 24** | Nil | Nil | Nil | *B. cereus* (3.0) |
| **Sample 25** | Nil | *S. epidermidis*(2.3) | Nil | Nil |
| **Sample 28** | Nil | *S. epidermidis*(3.0) | Nil | Nil |
| **Sample 30** | Nil | *S. epidermidis*(3.9) | Nil | Nil |

The highest level of contamination (3.7 log₁₀ CFU) was observed for *B. amyloliquefaciens* on the left side of the probe handle, followed by 3.5 log₁₀ CFU each for Propionibacterium spp. and Pseudomonas fulva on the top notch area of the probe (Table 1). There was 1.6 log₁₀ CFU count for both *Staphylococcus epidermidis* and *P. fulva* on the top notch area of the untreated probe (Table 1). Similarly for probe B, a variety of bacterial species were detected, including L. pneumophila, Staphylococcus, Pseudomonas, and Bacillus spp. Almost all sites on the probe were contaminated with different bacterial species. The highest microbial load (3.9 log₁₀) was observed for S. epidermidis on the left side of the probe’s top notch, followed by 3.8 log₁₀ for Pseudomonas koreensis on the right side of the groove and 3.6 log₁₀ for Bacillus cereus on the right-side of top notch (Table 1). The lowest bacterial counts 1.8 log₁₀ were detected for S. epidermidis in the right side of groove followed by 1.9 log₁₀ for S. capitis on left side of groove probe.

The rate of contamination of top notches of the probe was 57% (4/7), bottom notches 25% (2/8), right side of handle 43% (3/7), left side of handle 25% (2/8) before UVC treatment (Table 2). However, after UVC treatment, rate of contamination was 0% of all tested sites of probe (Table 2). Similarly, Probe B was also heavily contaminated with microbes before UVC treatment. All the sites shown (Table 2), were assessed for microbial presence and found contaminated on non-UVC treated sites. Rate of contamination of right side of edge 43% (3/7), left side of edge 38% (3/8), right side of groove 57% (4/7), left side of groove 25% (2/8). However, after UVC treatment, there was complete killing of microorganisms on each treated site, and no microbial growth was detected on any treated site The rate of contamination after UVC treatment on all the sites of probe was 0 % (Table 2).

**Table 2.** Sites of probes A and B showing bacterial contamination before and after UVC treatment.

| Location | Before UVC | % Contaminated Before UVC | After UVC | % Contaminated After UVC | P value |
| --- | --- | --- | --- | --- | --- |
| Probe A |  |  |  |  |  |
| Top Notch | 4/7 | 57 | 0/7 | 0 | <0.05 |
| Bottom Notch | 2/8 | 25 | 0/8 | 0 |  |
| Right Side of Handle | 3/7 | 43 | 0/7 | 0 |  |
| Left Side of Handle | 2/8 | 25 | 0/8 | 0 |  |
| Probe B |  |  |  |  |  |
| Right Side of Edge | 3/7 | 43 | 0/7 | 0 |  |
| Left Side of Edge | 3/8 | 38 | 0/8 | 0 |  |
| Right Side of Groove | 4/7 | 57 | 0/7 | 0 | <0.05 |

Some of the probe sheaths used for covering the probes were found to be contaminated. No microbial growth was detected on sheath 1. All other sheaths showed contamination with a range of bacterial species. Sheath 2 was contaminated with *S. epidermidis* (3.6 log₁₀ CFU). Sheath 3 showed contamination with *S.lugdunensis* and *Bacillus oceanisediminis,* with counts of 3.5 and 3.4 log₁₀ CFU, respectively. Sheath 4 was contaminated with *S.epidermidis*, exhibiting the highest bacterial load (4.3 log₁₀ CFU). Sheath 5 was found contaminated with multiple bacterial species, including *Nocardioides simplex* (4.2 log₁₀ CFU), *Corynebacterium aurimucosum* (3.9 log₁₀ CFU), and *S. epidermidis* (3.3 log₁₀ CFU) (Table 3).

**Table 3.** Numbers of bacteria identified on the sheaths (protective covering of probes).

| Covering | Bacterial type | Log<sub>10</sub> CFU |
| --- | --- | --- |
| Sheath 1 (from sample 10) | *Nil* | 0 |
| Sheath 2 (from sample 17) | *S. epidermidis* | 3.6 |
| Sheath 3 (from sample 21) | *S. lugdunensis* | 3.5 |
|  | *Bacillus oceanisediminis* | 3.4 |
| Sheath 4 (from sample 24) | *S. epidermidis* | 4.3 |
| Sheath 5 (from sample 29) | *Nocardioides simplex* | 4.2 |
|  | *Corynebacterium aurimucosum* | 3.9 |
|  | *S. epidermidis* | 3.3 |

The environments of both the clinical room (where samples were collected) and the disinfectant room (where samples were processed using the UVC device) were found to be contaminated throughout the study period. Overall, the clinical room tended to show higher levels of contamination than the disinfectant room on each day, with the types of bacteria found varying between days and rooms (Table 4).

**Table 4.** Numbers of bacteria identified in the environment where probes were sampled and processed.

| Collection day | Clinical/Examination Room |  | Disinfectant room |  |
| --- | --- | --- | --- | --- |
|  | Bacterial type | Numbers (Log<sub>10</sub> CFU) | Bacterial type | Numbers (Log<sub>10</sub> CFU) |
| Day 1 | *S. hominis* | 2.9 | *S. epidermidis* | 1.6 |
|  | *S. warneri* | 2.8 | *M. luteus* | 1.3 |
|  | *S. epidermidis* | 1.8 | *S. hominis* | 1.1 |
|  | *E. faecium* | 1.6 | *K. rhizophila* | 0.9 |
|  | *M. luteus* | 1.6 | *S. warneri* | 0.9 |
|  | *S. haemolyticus* | 1.2 | *E. faecium* | 0.8 |
|  | *Corynebacterium riegelii* | 1.1 |  |  |
| Day 2 | *K. rhizophila* | 1.9 | *S. hominis* | 1.1 |
|  | *B. cereus* | 1.9 | *S. pasteurii* | 0.9 |
|  | *S. hominis* | 1.7 | *S. haemolyticus* | 0.9 |
|  | *S. warneri* | 1.5 | *P. koreensis* | 0.6 |
|  |  |  | *S. epidermidis* | 0.6 |
| Day 3 | *S. haemolyticus* | 2.1 | *S. epidermidis* | 1.3 |
|  | *B. cereus* | 1.6 | *S. epidermidis* | 1.3 |
|  | *S. warneri* | 1.3 | *K. rhizophila* | 1.2 |
|  | *S. epidermidis* | 0.9 | *B. cereus* | 1 |
|  | *Demacoccus nishinomiyaensis* | 0.6 | *S. capitis* | 0.9 |
|  |  |  | *Streptococcus sanguinis* | 1.1 |
| Day 4 | *S. hominis* | 2.6 | *S. epidermidis* | 1.9 |
|  | *Corynebacterium aurimucosum* | 2.3 | *Dermatophilus congolensis* | 1.9 |
|  | *S. cohnii* | 1.9 | *M. luteus* | 1.6 |
|  | *Legionella pneumophila* | 1.8 | *Kocuria rhizophila* | 1.3 |
|  | *Micrococcus luteus* | 1.7 | *E. faecium* | 1.0 |
|  | *B. cereus* | 0.9 | *S. condiment* | 0.8 |
|  | *S. arlettae* | 0.6 | *S. lugdunensis* | 0.6 |

### In vitro testing

No spores germinated from the probes after disinfection with 4% bleach, indicating that this method was effective for decontaminating probes prior to use in the study as before inoculation of each repetition. In addition, no visible bacterial growth was observed in the sterile recovery broth after incubation for up to 24 h, confirming that the broth remained sterile. On other hand, visible growth was observed in the tube containing 1.0 mL of a 1 × 10⁷ spore dilution, demonstrating that the spores were viable. Spore suspensions adjusted to OD₆₂₀ values of 1- 3 yielded 1.5 × 10⁹ CFU/mL in hard water containing 5% horse serum.

The efficacy of UVC treatment in reducing microbial contamination on the complex probe sites of the handle, notch, and lens, was evaluated by comparing viable counts recovered from control and UVC-treated samples following five consecutive inoculations with *Bacillus* spores. Before UVC treatment, mean recoveries of 6.94±0.04, 7.12±0.15, and 6.82±0.12 log₁₀ CFU were observed on the handle, notch, and lens areas of probe A, respectively. Following UVC exposure, recoveries were reduced to 0.14±0.31, 0.20±0.45, and 0.00±0.00 log₁₀ CFU on the corresponding sites. Overall, UVC treatment achieved log₁₀ reductions of 6.80 on the handle, 6.92 on the notch, and 6.82 on the lens of probe A (Figure 2A). Similarly, for probe B, untreated control samples showed mean recoveries of 6.99±0.08, 7.11±0.40, and 6.85±0.08 log₁₀ CFU on the central bent side, left edge, and lens area, respectively. After UVC treatment, viable counts decreased to 0.20±0.45, 0.14±0.31, and 0.00±0.00 log₁₀ CFU on the same sites. Consequently, UVC exposure resulted in log₁₀ reductions of 6.79 on the central bent side, 6.97 on the left edge, and 6.85 on the lens area of probe B (Figure 2B).

**Figure 2.**
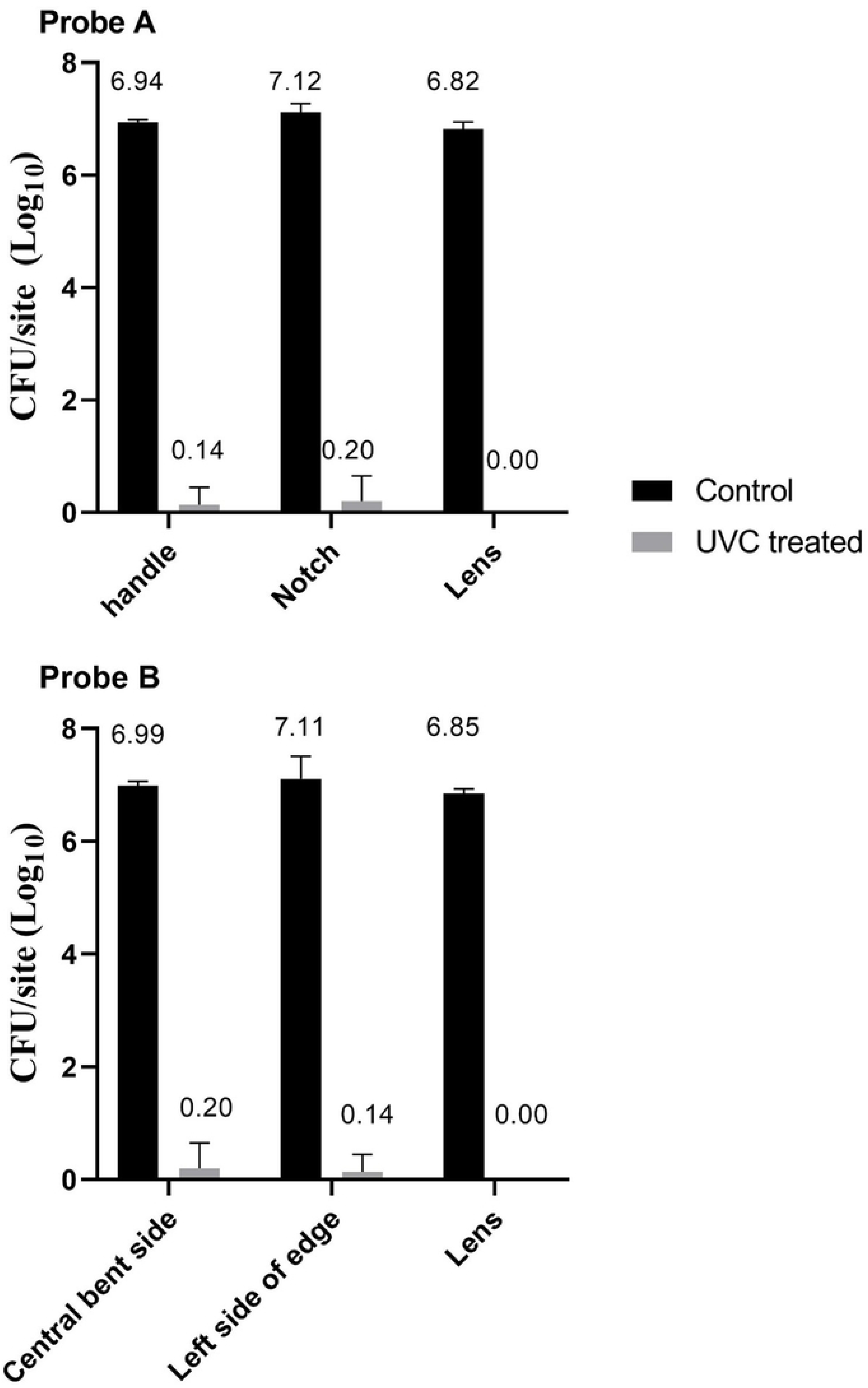
Effect of UVC treatment on microbial recovery from different sites of probes. Viable counts (log_10_ CFU/site) recovered from control and UVC-treated samples are shown for the handle, bottom notch and lens **(Probe A),** and central bent side, left side of edge and lens of **(Probe B).** Control sites exhibited high microbial loads > 6.5 log_10_ CFU/site, whereas UVC treated sites exhibited< O.Slog_10_ CFU/site. This indicated> 6 log_10_ reduction of bacillus spores was noted on complex sites of both probes.

Overall, UVC treatment resulted in complete inactivation of recoverable microorganisms on both probes and handle, bent areas, notch, edges and lenses, achieving reductions of approximately > 6 log₁₀ CFU/site when compared with untreated controls.

## Discussion

The present study demonstrates that probes can be extensively contaminated prior to disinfection, with multiple complex sites harbouring diverse and clinically relevant bacterial species. Before UVC exposure, contamination rates ranged from 25% to 57% across different probe regions including notches, handles, edges, and grooves. However, UVC LED system effectively reduced the microbial load by killing 100% on all these complex sites after treatment with standard cycle for 90 sec. The efficacy of UVC LED, was not affected by the microbial contaminants found in the environment where these probes have been disinfected.

Complex probe geometries such as depressed or irregular areas (notches and grooves and bent areas) provide favourable niches for microbial retention (13). In this study, two probes featuring such notches and grooves were examined to determine the presence of microorganisms following clinical use. These recessed areas provide sheltered sites that hinder routine cleaning and disinfection because of surface irregularities and limited accessibility. A diverse microbial community, including *Staphylococcus*, *Pseudomonas*, *Propionibacterium*, *Bacillus*, *Tsukamurella* spp., and *L. pneumophila*, was recovered from these hidden regions on untreated probes. Similar organisms have been previously reported on various probe surfaces (14). Moreover, the probe’s handle and can also become contaminated (15) where *Pseudomonas sp, Staphylococcus sp,* and *L. pneumophila* can be transferred from environmental surfaces or the skin of handling staff.

Two *Bacillus* species were found on the probes’ handles or notches (lens). This is similar to a previous study that also found *Bacillus* on probes (16). *Bacillus* sp. are spore forming bacteria and represent stable environmental reservoirs contributing to probe contamination. However, after treatment with UVC LED system no microbial growth was detected from any of these areas. This suggests that UV-C LEDs delivered UV-C light effectively across the entire probe surface using UVMESH technology, resulting in the elimination of microorganisms from complex and uneven probe surfaces.*Staphylococcus* species particularly *S. epidermidis*, *S. haemolyticus*, and *S. warneri*—were frequently recovered from complex probe sites. Notably, *S. epidermidis* reached levels of 3.9 log₁₀ CFU on the left side of the top notch of untreated probes. This location is known to be one of the most consistently contaminated areas of ultrasound probes. Previous studies also highlight the prevalence of *Staphylococcus* spp., especially *S. epidermidis*, on probe surfaces (6). Given that many of these organisms are common skin commensals (17), they can easily transfer from human skin to probes during handling. Minimising unnecessary contact after high-level disinfection is therefore essential to prevent reintroduction of microorganisms into these difficult-to-access probe areas.

Probe covers, although designed to act as protective barriers, can themselves become contaminated during ultrasound procedures through contact with mucous membranes, infectious sites, or open wounds. In the present study, four of five sheath samples were contaminated with *B. oceanisediminis*, *S. epidermidis*, *S. lugdunensis*, *Nocardioides simplex*, and *Corynebacterium aurimucosum*. Previous work has similarly shown that probe covers can transfer microorganisms to probes, serving as an additional contamination source (6). The high level of *S. epidermidis* detected (4.3 log₁₀ CFU) suggests compromised barrier integrity, improper handling, or environmental contamination during application or removal. Detection of multiple species on individual covers further indicates that the use of sheaths alone does not eliminate contamination risk and underscores the continued need for effective disinfection protocols even when protective barriers are used. Consistent with earlier recommendations, a new condom or probe cover should be applied for every patient and for each use of the instrument (18).

The detection of environmental and opportunistic pathogens such as *L. pneumophila* and *Pseudomonas* spp. indicates the potential for cross-contamination between clinical environments and medical devices. Environmental monitoring showed persistent background contamination in both the clinical and disinfectant rooms, with higher microbial loads in the clinical room but measurable contamination in both settings. Recurrent recovery of coagulase-negative *Staphylococci*, *Bacillus* spp., *Micrococcus* spp., and *Corynebacterium* spp. reflects ongoing environmental shedding and surface contamination. The microbial profile on the probes closely mirrored that of both rooms across the four sampling days. The dominance of *Staphylococcus* spp., including *S. hominis*, *S. epidermidis*, *S. haemolyticus*, *S. warneri*, and *S. cohnii*, paralleled their frequent detection on probe surfaces, consistent with their role as common skin commensals and environmental contaminants (17). Their presence in both settings strongly suggests transfer via handling or contact with contaminated surfaces. Similarly, *L. pneumophila* detected in the clinical room (1.8 log₁₀ CFU on Day 4) corresponded with *Legionella* species identified on the probe, indicating possible water or aerosol based contamination routes (19). The occurrence of *Pseudomonas* spp. in both environments *P. koreensis* in the disinfectant room and *P. fulva* on the probe supports moisture-associated transmission.

Higher overall contamination in the clinical room also aligned with the elevated microbial loads found on probes, particularly on frequently handled or structurally complex regions such as the handle and top notch. These areas likely promote microbial retention due to repeated manual contact and micro-topographical features that hinder effective cleaning.

In vitro validation confirmed the robustness of the experimental design. The absence of spore germination following 4% bleach treatment verified that decontamination was effective prior to inoculation experiments as demonstrated earlier (6). Recovery of viable *B. subtilis* spores and sterile control media further supported the accuracy of microbial reduction measurements. UVC treatment completely eliminated microorganisms across all probe sites. Under high-level *Bacillus* spore challenge conditions, UVC exposure achieved > 6 log₁₀ CFU reductions on all surface types. Such reductions meet accepted standards for high-level disinfection, even against highly resistant spore-formers. Minor residual counts (0.23–0.33 log₁₀ CFU) may reflect organisms near the detection limit rather than meaningful survival. *Bacillus* was selected because related species were isolated from used probes, spores are internationally recommended test organisms (EN12353, EN14561, AOAC 966.04), and they exhibit exceptional resistance to disinfection [15, 16]. Importantly, the efficacy of UVC was uniformly distributed across varied surface configurations, suggesting effective penetration and coverage despite geometric complexity of the probes.

The combined clinical and laboratory findings highlight two major implications. First, routine probe use in clinical environments can result in substantial and diverse microbial contamination, even when protective sheaths are utilized. Second, UVC disinfection provides rapid and highly effective microbial reduction, including against resistant spore-forming bacteria, making it a strong candidate for high-level disinfection of complex reusable probes.

This study has several strengths, including site-specific contamination mapping, identification of a broad spectrum of microorganisms, environmental monitoring, and rigorous spore-based challenge testing. However, certain limitations should be acknowledged for the recent study. Viral pathogens were not evaluated. Additionally, while environmental contamination was recognized, transmission dynamics were not directly investigated. Future research should explore the effectiveness of UVC against viral pathogens. Another, key limitation of some conventional disinfection approaches, where irregular surfaces can compromise microbial inactivation. However, the UV-C LED system combined with UVMESH technology similar to previous study (20), delivers comprehensive 360° UV-C coverage achieving microbial inactivation across complex probe surfaces in the current study. This supports the reliability and reproducibility of the UV-C disinfection system in real-world clinical environments. In conclusion, the findings demonstrate that reusable clinical probes are frequently contaminated with diverse microbial species under routine conditions. UVC treatment achieved consistent and substantial (> 6 log₁₀) microbial reductions across all tested sites, supporting its effectiveness as a high-level disinfection strategy for complex medical devices.

## Acknowledgement

The University of New South Wales School of Optometry and Vision Science, Faculty of Medicine and Health wishes to thank Connect IVF, Fertility Clinic located at 20 Bridge Street Sydney 2000, for helping in collection of the samples. We are also thankful to Anjaneyaswamy Ravipati (Swami) in this for helping in bacterial identification using MALDI TOF technique. **Conceptualization:** Muhammad Yasir.

Formal analysis: Muhammad Yasir

**Funding acquisition:** Mark D. P. Willcox. **Methodology:** Muhammad Yasir, Mark D. P. Willcox. **Project administration:** Mark D. P. Willcox.

**Resources:** Mark D. P. Willcox.

**Supervision:** Mark D. P. Willcox.

**Writing – original draft:** Muhammad Yasir.

**Writing – review & editing:** Muhammad Yasir, Mark D. P. Willcox.

